# Head direction cells use a head-referenced dual-axis updating rule in 3D space

**DOI:** 10.1101/2025.09.17.676760

**Authors:** M Williams, JS Street, N Burgess, KJ Jeffery

**Affiliations:** Department of Clinical and Movement Neurosciences, Institute of Neurology, University College London; Institute of Behavioural Neuroscience, University College London, where the experimental work was done; Department of Clinical and Experimental Epilepsy, UCL Queen Square Institute of Neurology, University College London; UCL Institute of Cognitive Neuroscience, University College London; School of Psychology & Neuroscience, University of Glasgow

## Abstract

Head direction (HD) cells, comprising a compass signal for the brain’s spatial map, may update their firing during 3D movement by using a ‘dual axis’ (DA) rule that summates head rotation around a local axis and local-axis rotation around the gravity axis. To test for operation of this rule we recorded HD cell activity from rats exploring a hemispherical surface. We assessed HD cell tuning curves in either the standard horizontal reference frame or in two reference frames governed by a DA rule, with the local axis referenced to the head (DAH) or local surface (DAS). Tuning curves were best when plotted in the DAH reference frame. This confirms that the cells use a head-referenced DA rule and indicates that the relationship of the head pose to gravity in two planes, one egocentric (head-plane) and one allocentric (earth-horizontal plane), forms a component of the inputs to the HD system.

## Introduction

Spatial cognition in large-scale space requires a stable internal compass, which is implemented by the head direction (HD) neurons found in several brain regions. HD cells signal the azimuth (horizontal facing direction) of the head, or the “nose vector,” with directionally oriented tuning curves, firing maximally when the nose vector aligns with the cell’s preferred firing direction (PFD).^1^ For a given cell, firing falls off smoothly as the nose vector’s angular distance from the PFD, called alpha (*α*), increases, falling to zero at approximately *α* = +/-45 degrees. Network activity is organized as a hypothetical “ring attractor,” ^2–4^ updating in response to the left-right head-turns that dynamically vary *α*.

A difficulty with this scheme is that during 3D movement, such as on a sphere (Fig. 1a), a ring attractor network with a purely left-right-based (“yaw”) updating rule would accumulate so-called Berry-Hannay errors^5^ due to occult changes in azimuth arising from non-horizontal 3D rotations (Supp. Fig. 1a). However, HD cells do maintain a stable PFD on a sphere (see example cells in Fig. 1b and Supp. Fig. 5), indicating an ability to correct these errors. This might indicate pure reliance on external directional environmental cues, but such reliance would be vulnerable to losing sight of these cues: e.g., when looking elsewhere, or in the dark. On the horizontal plane, this vulnerability is mitigated by pairing visual orienting with self-motion tracking (mainly vestibular-based) so as to keep the signal stable in between visual “fixes.” ^6,7^ It has been suggested that this scheme could still work in 3D if HD cells use a so-called “dual axis” (DA) rule which takes into account not just the rotation of the head around its own axis (or an axis defined by the local surface; see below), but also the rotation of that axis around the gravity axis.^8–10^ These rotations would be summed to yield a local PFD, adjusting with rotation of the head’s dorso-ventral (DV) axis through space (Fig. 1c) such that on returning to a horizontal pose, the local and global PFDs have retained alignment. This proposed modification to the ring attractor network architecture would allow effective navigation in 3D using only a 2D representational substrate, even during absence of external vision, thus maintaining global directional orientation with minimal neural resource use.

**Figure 1:**
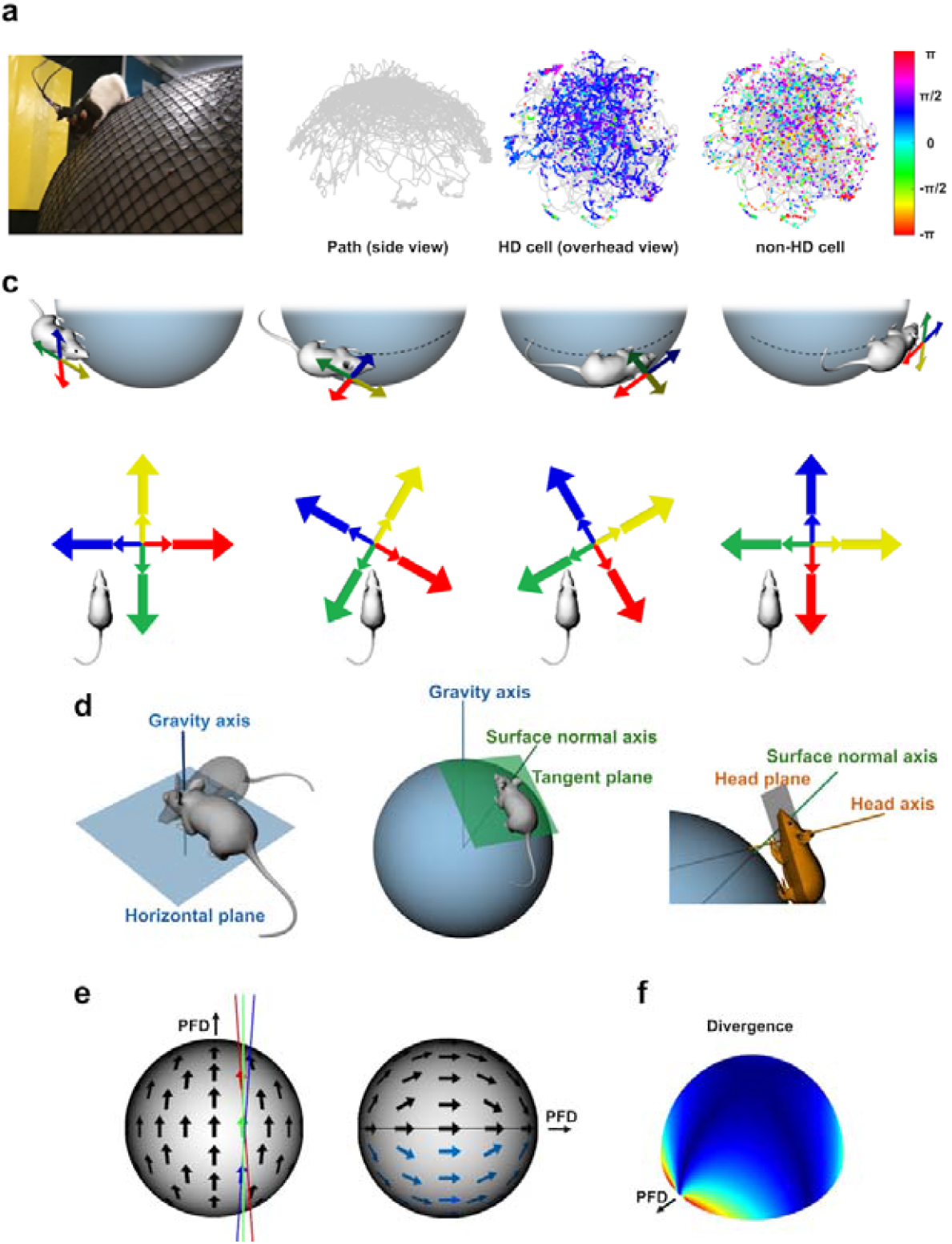
Dual axis updating of HD cells (a) Rat exploring a hemispherical surface (frame from video^8^). (b) Position and spike data from a single sphere recording session. The path of the animal is shown in gray, and spikes are shown as dots color-coded by azimuth (see color bar). Left-right: Path from side view, example HD cell, and example non-HD cell. (c) Correcting for rotations of the head around the gravity axis (See Supp. Fig. 1 for more details). By a DA rule, as the rat progresses around the sphere its local reference frame rotates opposite to the rotation direction of travel to maintain alignment with the global frame. (d) The three orientational reference frames (each described by an axis and its plane) examined in this study. (e) Vector fields from a hypothetical HD cell showing the horizontal pointing direction (global azimuth) of the cell’s PFD as seen from above (left) or the side (right). The sphere represents either an allocentric reference frame (for the DAH model) or an egocentric, head-centered reference frame (for the DAS model). The colored lines on the overhead view show the subtle divergence from parallel under the DA rules. (f) Theoretical divergence of local PFDs from the global PFD, as a function of position on the sphere (hot colors = greatest).

Support for use of the DA rule has been reported by Page et al.,^8^ Laurens and Angelaki,^9,10^ Shinder and Taube^11^ and LaChance et al^12^. These experiments were conducted under restricted conditions, either when the animals were confined to a plane or a small number of planes, or when they were restrained and passively rotated. These studies provided evidence consistent with operation of a DA rule, including preserved directional tuning on a vertical surface, updating when the head rotated in its own plane, and rotation of the local PFD when the animal’s DV axis rotated around a vertical axis such as a corner. However they did not determine whether the DA rule is continuously used during unrestricted navigation over a continuously curved surface, which is the more naturalistic condition.

Additionally, previous HD cell experiments with overhead cameras have referenced the signal to the horizontal plane: whether the more appropriate reference frame is the plane itself or the head of the animal has not been experimentally disambiguated. Either option is plausible, given the existence of environment-referenced and self-referenced encoding schemes throughout the spatial system^.13^ One potential reason for the system to use an environmental reference frame is that a directional signal that remained stable within a local bounded frame might be easier for grid and place cells to interpret in service of their own spatial computations. However, Shinder and Taube^11^ have argued in favor of the head-referenced frame, given the plentiful vestibular inputs available to this system.

In the present experiment we thus sought to answer two outstanding questions: (i) Does the DA rule operate in free movement over a curved surface, even when distal visual cues are visible and head-axis orientation changes smoothly rather than abruptly? And (ii) if so, is the second axis is supplied by the environment or by the azimuth of the head axis? To answer these questions we recorded HD cells from postsubiculum (PoS) and dorsal anterior thalamus (ADN) in rats exploring a hemispherical surface (see Supp. Movie).^14^ We refer to three rotational reference frames, each defined by an axis of rotation plus its corresponding orthogonal plane (Fig. 1d), as follows: (i) the gravity axis (GA) plus horizontal plane, (ii) the environment’s surface-normal axis (SA) plus its tangent plane, and (iii) the dorso-ventral head axis (HA) plus the transverse plane of the head. The DA rule using the SA axis is referred to as DAS, and using the HA is DAH.

We thus investigated whether the PFDs of the neurons would be best described by a purely horizontally referenced updating rule (GA) describing global azimuth only, or better described by the DAS or DAH, a transformation (Fig. 1e) that should be greatest at the azimuthal poles defined by the PFD (Fig. 1f). We compared the data with a null reference frame in which only rotations about the head axis (HA) were tracked. We show here that when the data are plotted in a dual axis reference frame the tuning curves are taller and sharper (more precise) and are better fitted by the von Mises function that describes head direction cell tuning curves. This improvement is much stronger for DAH than DAS. These findings support the dual-axis hypothesis and confirm that the local axis is the head rather than the environment surface.

## Results

To investigate whether a dual axis rule better describes HD firing in 3D than the GA rule that applies in the horizontal plane, we recorded 2270 single units (436 HD cells), using tetrodes, in 10 rats from either anterodorsal nucleus of the thalamus (ADN, n = 5) or postsubiculum (PoS, n = 5) while the animals freely foraged over the upper surface of a 1-m diameter sphere. Position in 3D coordinates was determined by tracking a head-mounted LED with 5 cameras, and 3D orientation of their heads using an inertial measurement unit (IMU) that was calibrated with a visual ground truth (Supp Fig. 2).

The animals did not sample all directions uniformly over the hemisphere (Figs 1a and Supp. 3a), preventing assessment of full tuning curves everywhere over the sphere surface. Instead, we adopted a model-fitting approach based on inferring the “local *α*” at each moment in a local plane relative to a local PFD. Local *α* was computed according to four different models, and used to generate modified tuning curve that were compared with the original ones. The analysis pipeline is described in detail in the Methods and shown in Supp. Fig. 4c. Briefly, it runs as follows: first, we smoothed the spike trains, binned in time bins of 20 ms and then directionally binned the data in 6-deg. bins according to the corresponding nose vector projected onto the horizontal plane. This yielded “global azimuth” tuning curve histograms, from which we calculated peak firing rates and Rayleigh vectors. We then derived a model to describe these tuning curves using a simplified version of the von Mises equation for HD cells (Equ. 1), relating firing rate to the nose vector’s *α* as follows:

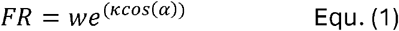

Given a set of nose vectors and firing rates, this equation can be used in a search process to discover the parameters of *PFD, κ* (kappa; tuning curve sharpness) and *w* (weight; equivalent to peak) for a given cell’s trial data. After applying this procedure to the global azimuth tuning curves we selected, from these, the subset of cells (n = 436) having *κ* > 0.5. This unbiased selection produced results that agreed well with visual assessment of tuning curve quality, and the cells had typical tuning curve properties in a standard horizontal reference frame (Supp. Table 1 and Supp. Fig. 5, left-hand columns). Using the derived model parameters, we generated for each cell a model tuning curve, and generated an initial model fit to the actual tuning curve histograms using Pearson’s correlation (R). This procedure yielded the GA model.

We then transformed the data by hypothesizing that cells are better described with reference not to global PFD but to a local PFD, this being the global PFD modulated by a DA rule. We inferred the local *α* at each moment by computing and summing two rotations: (i) the yaw (left-right) rotation of the head in either its own plane (DAH) or the tangent plane of the sphere (DAS) relative to vertical (the GA, projected onto the plane), and (ii) the yaw rotation of either the SA or HA in the horizontal plane, relative to the point on the circle antipodal to the global PFD (that is, PFD + *π* radians). This was chosen as the reference point because it is where *α* is known, from prior work,^15^ to be zero when the nose points directly upwards. As a null control, we computed *α* in the egocentric reference plane of the rat’s head around the head axis (HA; a single-axis purely egocentric rule).

Using the resultant set of local *α’s* we then re-plotted the tuning curves to determine the new peak rates and Rayleigh vectors. We then used the same model search process as we had used previously to re-derive new κ and *w* model parameters, and fitted the new model to these.

An illustration of this procedure (see Online Methods for details) is shown for the example cell in Fig. 2a, and a broader selection in Supp. Fig. 5. In the plots, the tuning curves have been created using *α*’s from the four transformations. With the two DA transformations the tuning is sharpened, both visibly and as reflected in the peak-rate and κ values. This effect is more pronounced for DAH than DAS.

**Figure 2:**
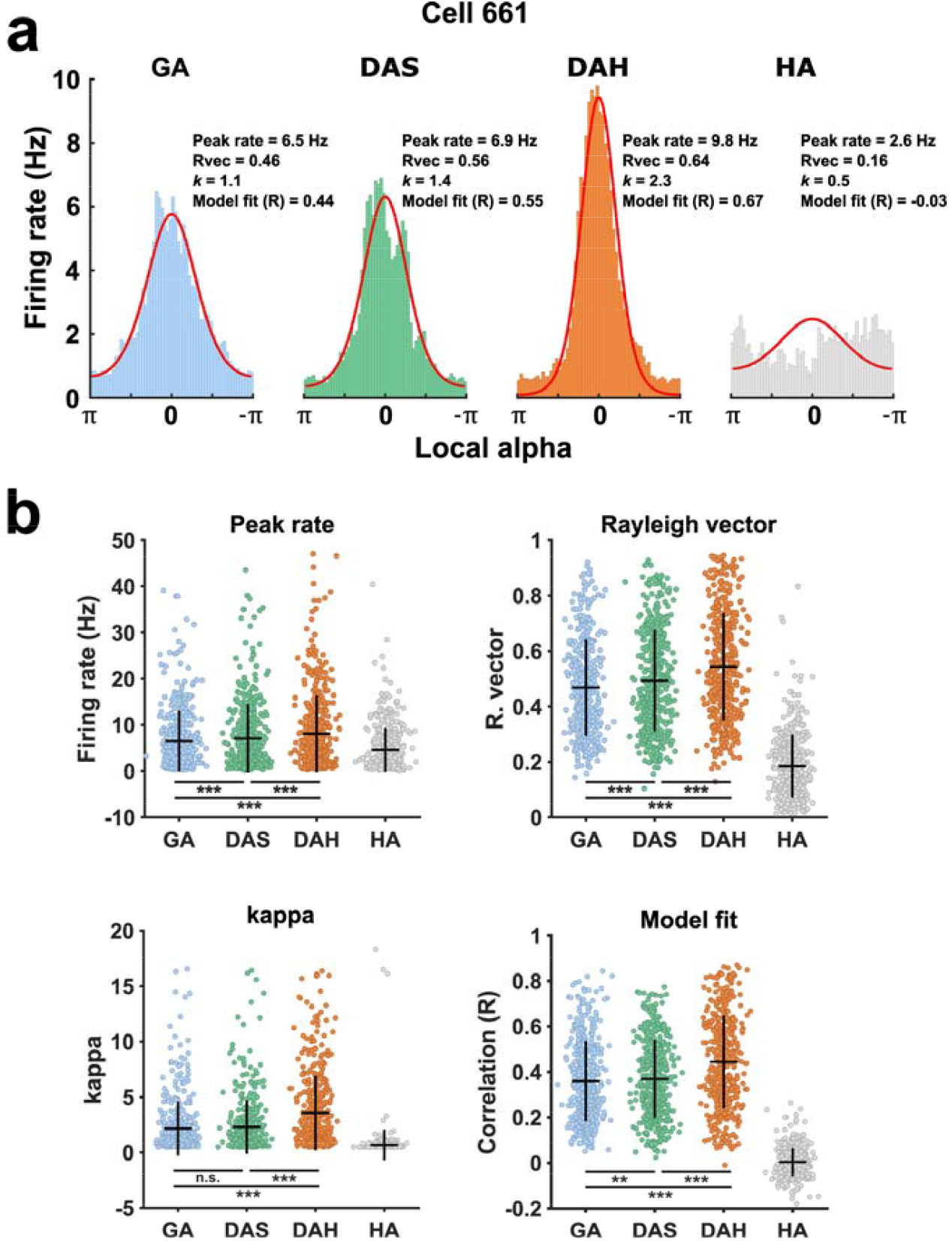
Sharpening of tuning curves with change in reference frame. (a) Model fit procedure. For this single ADN cell, four tuning curves (directional firing rate histograms have been generated in, respectively, the standard gravity-axis reference frame (GA), surface-axis referenced dual-axis frame (DAS), head-referenced dual-axis frame (DAH) or head-referenced only frame (HA). The DAH tuning curve was tallest (Peak rate) and sharpest (kappa) with the greatest directionality (Rayleigh vector) and best von Mises model fit (red line). For further examples see Supp. Fig. 5. (b) Tuning curve and model fit parameters for all HD cells (n = 436; ** = p < 0.01, *** = p < 0.001).

It is apparent by visual inspection that the majority of tuning curves were best (tallest and narrowest) when plotted under the DAH model. To quantify this we first used the Pearson’s R model fit parameter (Supp. Fig. 4c) as the measure of a cell’s best model and counted the cells for which each model was the best fit (Supp. Table 2). Cell counts differed markedly across models (χ^2^(3) = 916.6, p < 10^−16^). Standardized residuals showed that DAH was the best model for significantly more cells.

We then used linear mixed-effects models to examine the effect of the four models on four response variables (peak firing rate, Rayleigh vector length, κ, and model fit). All models were fitted using maximum likelihood estimation, with model type as a fixed effect and random intercepts for rat and for cell nested within rat. Analyses were conducted using MATLAB’s *fitlme* function. For each outcome, we assessed the omnibus fixed effect of model and then conducted planned contrasts reflecting a priori hypotheses: (i) DAH vs each alternative model, and (ii) DAS vs GA. The results are summarized in Supp. Table 3. *Model* had a strong fixed effect for all measures. Estimated marginal means consistently ranked DAH highest, and planned contrasts confirmed the a priori hypothesis that DAH exceeded DAS, GA, and HA for every measure. The secondary hypothesis that DAS outperformed GA was supported for peak rate and Rayleigh vector length but not for κ or the model fit metric. Thus, DAH was a stronger model than DAS. The pairwise comparisons for the GA vs. other models illustrate the robustness of the effect across the cell population (Fig. 3) and the superiority of DAH over DAS (Fig. 3 rightmost column).

**Figure 3:**
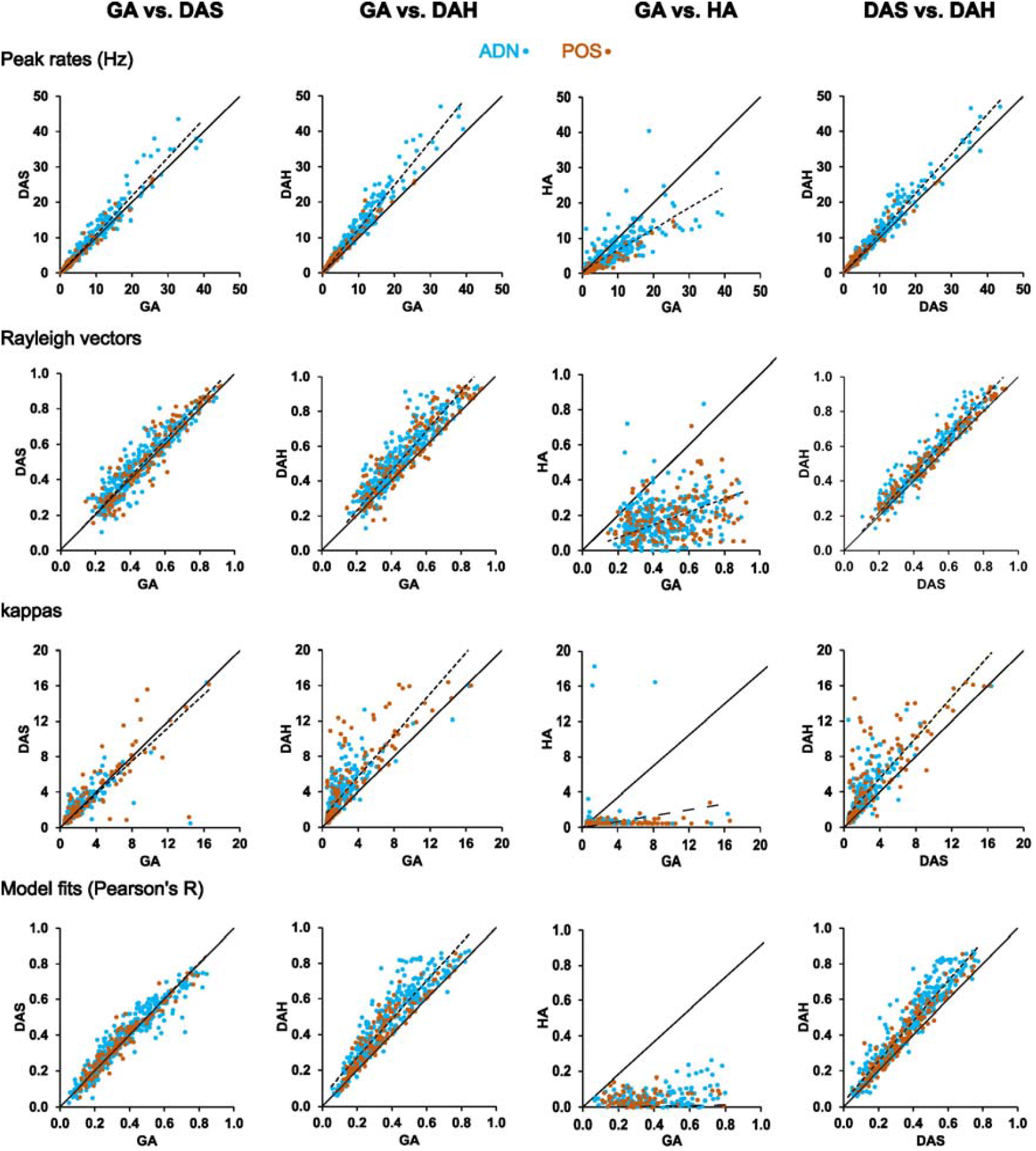
Model comparisons for the 436 HD cells. Pairwise comparisons between GA and DAS, DAH and HA models, plus a direct DAS-DAH comparison, for the four parameters from the tuning curve histograms. The model fit represents the Pearson’s R of the von Mises function fitted to the re-framed tuning curves. Solid black line shows y = x; dotted black line shows a linear fit to the actual data (ADN and POS combined). Note that the DAH model consistently outperforms the others (points lie above the y = x line) for almost all cells.

Operation of a DA rule requires detection of the tilt of the head relative to gravity, raising the possibility of finding cells tuned to this parameter in our data set, as Angelaki et al. reported in mice.^10^ This could have been evident as cells tuned to a particular elevation, or cells with tuning curves in the HA reference frame. We first looked at firing rate as a function of pure elevation (angle of the nose above or below horizontal). Behavioral sampling of elevation angles was restricted, (Supp. Fig. 6), with angles distributed around a peak of −0.52 radians, consistent with the observed preference of the animals for circumferential and downwards travel. We found no clear evidence for cells tuned to elevation: a plot of firing rates vs. elevation showed increased variance at the higher elevations (Supp. Fig. 6b) but this was likely due to the dwell-time undersampling. Neither visual inspection nor statistical analysis (not reported) found evidence of a systematic elevation-rate relationship, although this cannot be ruled out given the limited pitch sampling. However the observation is consistent with previous work by Shinder and Taube ^11^. We also looked for tuning in the HA reference frame. In our data (from ADN and POS combined) none of the 436 neurons had the HA reference frame as their dominant model.

## Discussion

We investigated how HD cells, which function to signal global head azimuth, solve the problem of stably updating their network activity during movement in three dimensions. We tested the hypothesis that they do so using a dual axis (DA) rule that links two reference frames, defined respectively by a local (head or local surface) axis and a global (gravity) axis. We tested this hypothesis against two competing hypotheses: that updating is purely egocentric (using the head axis only) or purely allocentric (using the gravity axis only). We found that in rats exploring the upper surface of a sphere, HD cell responses were best described by a DA model in which the local axis is the head and not the environment surface. The implications of this finding are outlined below.

### Tuning curve description in multiple reference frames

Although we have here described the head rotations in two separate (albeit linked) reference frames, mathematically the two summed rotations equate to a rotation around a single tilted axis; hence “tilted azimuth” in Laurens and Angelaki’s formulation ^9^ or “tilt-compensated compass” (TCC) in engineering terminology. Furthermore, as discussed later, although HD processing can be described in terms of dynamic updating using detection of rotation (a velocity signal), a DA scheme can also account for static establishment of the signal (“setting” and “resetting”) occurring independently of velocity and previous network state.

To describe the DA updating rule mathematically, we can re-formulate the HD cell von Mises function to account for the second axis, as follows:

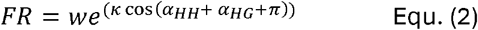

where *α*_*HH*_ is the alpha of the head rotation in the head plane around the HA (referenced to the projection of the GA onto the head plane) and *α*_*HG*_ is rotation of the HA in the horizontal plane around the GA.

Below, we discuss the adaptive advantages of the DA scheme, before considering possible sources of the two rotational signals onto the HD ring attractor network.

### Implications of a planar HD signal

The DA scheme yields a planar signal, as contrasted with a fully volumetric 3D signal which would encode the nose vector in all three rotational planes. This lower dimensional representational format is therefore unable to signal every possible head orientation in 3D space. In the case of rats, the missing signal is the amount of roll around the nose axis, which does not modulate the signal for rotations of at least +/-90 deg. ^11^

Why would evolution sacrifice this third degree of freedom? The likely answer is that because navigation over the surface of the planet is greatly weighted towards horizontal travel, it is inefficient to expend neural resource on representing directions that are rarely visited. The DA scheme allows the brain to maintain a continuous estimate of the global azimuth of the head in the horizontal plane using relatively few neurons that are in constant use. Since most animals locomote with the head held horizontal (or close to it)^16^, it makes adaptive sense to concentrate resources on representing the most behaviorally relevant plane.

Given that HD cells appear to form a planar compass, a planar variant that we considered is a purely horizontal one that detects and reports heading as global azimuth only, this being the horizontal projection of the nose vector. When we expressed HD cell firing in terms of the GA rule then tuning curves were well-formed (Supp. Fig. 5), which is consistent with a purely horizontal compass. However, they improved further when we expressed firing in terms of a local azimuth by implementing the DAH rule, which subtly deforms the HD cell vector field. Since tuning curves are usually plotted in horizontal coordinates only, and given that rats do not keep their heads completely horizontal during locomotion, it seems likely that HD cell intrinsic tuning curve widths have been overestimated in the past. For practical purposes, however, the signals are similar, so why should evolution have settled on the DAH rule? The answer may be that it is simply easier to compute. As the nose vector approaches vertical then its horizontal projection becomes vanishingly small, and it may be hard to stably drive the attractor with such a small signal; maintaining a local azimuth signal and then just modulating this is perhaps computationally easier. Possible mechanisms for the modulation are discussed in the next section.

A problem with any planar signal for an animal moving in three dimensions arises from the so-called Hairy Ball theorem, which asserts that on any closed surface having no holes, one cannot assign a smooth vector field everywhere without creating at least one singularity where the vectors collide. For the nose vector field over the surface of a sphere (Supp. Fig. 8a), this means that there must be two places where the attractor signal suddenly needs to jump to the far side of the ring: for example, when the animal crosses pitch-wise over the equator and its nose vector reverses. Another problem is that as the animal explores the under-surface of the sphere, the relationship between head-turns and allocentric direction reverses (Supp. Fig. 8b): where normally a head-turn to the right would shift the signal clockwise around the global azimuth, in inversion it shifts it counterclockwise. These two issues – singularities, and reversal of the egocentric/allocentric rotational relationship – might pose difficult problems for the HD ring attractor system. In support of this limitation, studies have suggested that the directionality of the HD signal breaks down in inversion^11,17^ along with the ability to navigate^18^. However, these experiments were conducted in rats, which have difficulty locomoting upside down due to their weight, and were perhaps anxious and distracted, and inattentive to spatial cues. A study of bats, which are comfortable roosting upside down, found that the directionality of the HD signal instead reversed in inversion: HD cells continued to fire when the animal’s nose vector crossed the equator, so that a “North”-indicating cell was now firing to the “South” ^19^. While this avoided the singularity problem for this pitch action, it would have created one had the animals instead rolled into inversion (roll was not assessed in this experiment). Overall, the picture is incomplete and further research is needed to determine how different species integrate directional signals throughout the full gamut of 3D movements, and how the signal feeds into navigational planning.

What might a planar HD signal imply for the processing of spatial cells – most prominently place cells, grid cells and border cells – that use the HD cell signal to orient their firing patterns? The implication is that the neural cognitive map is not truly volumetric: a point that has emerged from prior studies from experiments in 3D space. Although place cells pack a volumetric space with globular firing fields ^20,21^, grid cells do not form regular arrays of grid fields ^22,23^, which suggests that they do not make use of a fully 3D compass. However although the DA encoding scheme enables a planar compass to be efficiently deployed during 3D travel, the issue of singularities and inversion described above suggests that grid-cell grids, at least, might be predicted to break down in inversion. Studies of spatial encoding as animals explore in inversion as well as upright will be needed to understand how the system copes with these representational constraints. It is also interesting to speculate the constraints these processes place on other types of non-spatial cognition that have been posited to use the spatial system^24^.

### Network updating vs resetting in a dual-axis coding scheme

How are the two axes of the DA rule processed and integrated by the HD network? The HD signal, as studied in a purely horizontal reference frame, is well established to combine two elements: a dynamic updating signal that is linked to the network’s own activity at the preceding moment via a velocity signal, and a static resetting signal that comes from the environmental landmarks^2,3^. The updating signals come primarily from the vestibular system, although there is a contribution from other self-motion cues such as optic flow, proprioception and motor efference copy.^7,25^ Vestibular signals comprise rotational signals from the lateral (also called horizontal) semicircular canals, which detect left/right head movements and pass this information as a velocity signal to the central vestibular nuclei^26^. From here, this signal passes via the dorsal tegmental nuclei in the brainstem to the lateral mammillary nuclei and thence to the rest of the head direction system^27^. This velocity signal causes activity to shift around the head direction ring attractor at the appropriate rate, via a mechanism that has not yet been elucidated.

The environmental landmarks that provide static resetting information are usually (but not always) visual ^28^, and the signals are routed via primary visual cortex ^29^ into the head direction network. This is an instantaneous process that does not depend on preceding network state, and functions both to keep the network stably aligned across separate visits to the environment and between neighboring environments, ^6,30,31^ and also to provide moment-by-moment stabilization of the signal during a given visit. ^29,32^

How might this two-inputs system work for the two axes of the DA encoding scheme? We consider here the four elements (the two axes, and updating vs. resetting) as follows: (i) dynamic rotations around the egocentric head axis, (ii) static angle relative to gravity around the head axis, (iii) dynamic rotations of the head axis around the gravity axis, and (iv) static angle of the head axis relative to landmarks around the gravity axis.

- Dynamic rotations around the head axis. Even if the head is not horizontal, the head-axis velocity signal can be generated using the lateral semicircular canals, which still report left/right head-turns. Taube and colleagues have shown that the HD system has a bias for using information generated by head-turns around the head axis;^11^ (see Taube and Shinder^33^ for discussion).
- Static angle around the head axis. For the static resetting part of the process, which requires determination of environmental variables and is independent of prior network state, some kind of fixed reference direction is needed, against which the angle can be computed. This presumably would be the gravity vector, which is always present. Thus, setting/resetting could take place by detecting the instantaneous angle between nose vector and gravity. Consistent with this possibility, Angelaki et al. have reported neurons sensitive to the static tilt of the head in the anterior thalamus of macaques^34^ and mice.^10^ It is worth noting however that we did not see such responses in ADN, which should have manifested as neurons with tuning curves in the HA reference frame (that is, a given tilt of the head in the head-plane irrespective of head-axis azimuth). Nevertheless, tilt signals are present in other parts of the vestibular system. ^35,36^
- Dynamic rotations of the head axis around the gravity axis. Day and Fitzpatrick ^37^ showed that a neural process computes the vector dot product between the vestibular vector of head rotation around the head axis, and the gravitational unit vector, which yields horizontal rotational velocity. This requires a combination of the otolith organs to provide the gravity vector and the semicircular canals to supply the rotational signals. This integration may occur in the cerebellum: Yakusheva et al. ^38^ found that Purkinje cells recorded from the macaque cerebellar vermis encode a semicircular canal-driven signal that reflects the earth-horizontal component of rotation.
- Static head-axis angle relative to landmarks around the gravity axis. Can this be computed, and if so how? This would necessarily involve the environmental landmark panorama, of which many (albeit not all) landmarks would still be visible from a non-horizontal pose. For example, on a vertical surface with the nose pointing up then although the landmarks straight ahead of the animal are no longer visible, those to the left and right are (Supp. Fig. 7). These can in principle input to the ring attractor in the usual way, assuming continued object recognition given the now-tilted viewing angle. When the animal yaws its head around the head axis, the left and right landmarks now change their perceived angles; for a rightwards head turn the right-hand landmark is now “up” in retinal coordinates and the left landmark “down”. These angles could in principle be used to drive the ring attractor at its appropriate location, perhaps supported by the otolith signal that confirms the degree of head tilt relative to gravity.

Collectively, then it seems that the setting/resetting/updating processes determined for horizontal HD cell activity can feasibly work in a similar way for two axes. Exactly where these signals all converge, and how they integrate at a cellular level, is still a matter for investigation.

## Conclusion

This experiment has confirmed the hypothesis that HD cells combine head rotations in two independent reference frames, one allocentric, organized around the gravity axis, and one egocentric, organized around the head axis. Gravity is important in both frames: it provides a rotation axis for the allocentric reference frame and a fixed reference point for the egocentric reference frame. The adaptive consequence of this dual axis scheme is that the facing direction of the head can be continuously related to the Earth-horizontal frame using just a one-dimensional ring attractor, without accruing errors during 3D movement. Exactly where and how these rotational and gravity signals are combined and fed into the ring attractor network remains to be determined, as does the behavior of the network when it encounters the singularities that occur with some trajectories into inversion. This system is a paradigmatic example of dimensionality reduction, which is emerging as a brain organizing principle and may have implications for other types of cognition beyond simply spatial.

## Methods

### Experimental procedures

#### Animals and Care

Subjects were 10 adult male Lister-Hooded rats, obtained from Charles River, housed in a 11hr/11hr day/night cycle, with 1hr of dawn and 1hr of dusk. Pre-surgery, animals were housed in groups of 2-6 in a large cage (dimensions ≈2m x 2m x 2m). They were provided with apparatus to encourage 3D movement, including boxes, ledges, ropes, wooden balls, and spherical and volumetric climbing apparatus. Post-implantation they were housed singly but in clear-topped cages so they could see each other. All experiments were performed under UCL’s establishment licence and policies, in accordance with the Animal (Scientific Procedures) Act (1986).

#### Surgery

Surgery was performed once animals reached 350g. Animals were anesthetized with isoflurane and implanted with four tetrodes attached to a microdrive using standard stereotaxic procedures. Coordinates used for ADN were (in mm for AP, ML and DV respectively) bregma −1.8, 1.3, −4.0, and for PoS −7.5, 3.3, −2.0. After the tetrodes had reached the appropriate coordinates, the outer cannula of the drive was lowered to the brain surface, sealed with petroleum jelly, and fixed in place with dental cement. Once the drive was fully secure, the wire to the ground pin was soldered to the ground wire of the drive, and the animal was removed from the stereotaxic frame. Post-operatively, animals recovered in a heated recovery box until mobile. Analgesia was given for 3 days in the form of meloxicam (0.2mg/Kg) in condensed milk, and wet high-protein mash. Animals were closely monitored for 7 days, after which experiments began.

#### Apparatus

The screening arena consisted of a 1.2m x 1.2m open topped wooden box, with 60cm walls painted grey, a black antistatic polyvinyl floor, and a white cue card on one wall. The arena was set in a room with ample distal cues, including the recording PC, and the experimenter themselves.

In the experimental room, a 1m diameter spherical recording arena was situated in the centre of the room, surrounded by clear distal landmark cues on three walls, with the neural recording setup, PC, and experimenter behind a curtain on one side of the room to avoid distracting the rat. The sphere was constructed from paper mâché with the bottom 1⁄4 cut off to form a flat base. It was painted with non-reflective grey paint, and covered with a plastic mesh to allow animals to grip the surface. A 15cm ledge around the bottom prevented animals falling to the floor, and the whole apparatus was placed on a 40cm high trolley to prevent escape. Since the animals never strayed below the equator, we refer to it as the hemisphere. The cable of the neural recording system was suspended with elasticated thread with a system of pulleys to increase travel distance and allow the animal to move over the hemisphere unimpeded. Recordings were 20-30 minutes, depending on animal willingness to forage.

#### Recording Protocol

For hemisphere sessions, the headstage was plugged into the microdrive attached to the rat’s head, while the rat was distracted with a cereal reward. Once plugged in, the rat was placed on the hemisphere, where he foraged for malt paste until satisfied; usually up to 30 minutes. Malt paste was manually reapplied approximately every 5 minutes, primarily around the sides of the hemisphere to motivate exploration on the steeper surface. Except for when re-baiting the hemisphere the experimenter remained at the PC, behind a curtain, in order to monitor orientation, position, and neural recording quality. If the wire to the headstage became twisted, malt paste was used as bait to persuade the rat to turn back the other direction to untwist the cable. After recording concluded, the rat was picked up, given another reward, the headstage disconnected.

#### Screening and recording

Animals were screened for HD cells daily. Recordings were performed using an Axona recording system. Tetrodes were advanced by 50μm each day after recording. Screening sessions lasted 6 minutes each, after which the experimenter performed a brief assessment of cell clusters and HDC presence in TINT (Axona). If HD cells were present, the animal was moved on to the main recording session.

#### Positional Tracking

On the hemisphere, 3D position was recorded at 50Hz using a 5-camera setup (see ^21^. A 1Hz TTL pulse from the recording system drove an LED in the frame of each camera, which was used to synchronize the 5 analogue video streams offline. Camera feeds were digitized and adjusted for lens distortion, and 3D reconstructions of the position of a LED on the headstage were made using the MATLAB function triangulateMultiview.

#### Orientation Tracking

3D Head orientation was recorded on the hemisphere using a FSM300 (Hillcrest) inertial measurement unit (IMU). The IMU was mounted to the headstage such that it was level with the head (as defined by lambda and bregma), with the Y axis aligned with the nose, the X axis in the direction of the right ear, and the Z axis aligned with the dorsoventral axis of the animal. The same 1Hz TTL pulse used to synchronise the cameras was used for the IMU. The output of the IMU includes an accuracy metric, 0 being lowest and 3 highest. Data for which the accuracy metric was below 3 (meaning estimated error > 2°) was not used in analysis – these data typically amounted to 0-5% of a session’s data.

By recording HDCs and azimuthal nose direction in two sessions, one using the traditional LED method, one using the IMU, we confirmed that the IMU was able to provide accurate enough readings to identify HDCs (Supp Fig. 2). Simultaneous recording of the orientation of the IMU and LED apparatus, rotated by the experimenter, confirmed the accuracy of the IMU.

#### Histology

At the end of recordings, once no new HDCs had been recorded for 7 days, animals were culled and brains extracted. Anaesthesia was induced with 5% isoflurane, then animals were given an intraperitoneal (IP) overdose of pento-barbital (Pentoject). While the pentobarbital took effect, lesions were made at the tips of two tetrodes to aid visualisation post-mortem. After breathing and reflexes had ceased, animals were perfused with phosphate buffered saline (PBS), then 10% formalin in order to fix brain tissue. The brain was carefully extracted from the skull, placed in 10% formalin and refrigerated for at least 3 days. Brains were sliced into coronal sections of width 50μm, and mounted on glass slides. After drying, sections were stained with cresyl violet, washed with water and ethanol, and coverslipped. Sections were imaged using a bright-field microscope (Leica DMi8), and tetrode placement was assessed manually by the experimenter using a rat brain atlas (Paxinos and Watson, 2013).

### Neural data analysis

#### Clustering Single Units

To identify putative individual neurons, single unit data were clustered offline using Tint (Axona Ltd), which performs automated clustering based on principal component analysis (PCA) and an expectation-maximisation algorithm (Klusta-kwik ^39^. The resultant clusters were then manually curated. Clusters were excluded based on violation of a 2ms refractory period, or if they contained < 100 spikes. Once HD cells were identified (see below), the data were further refined to remove possible duplicates by identifying cells recorded in the same session from the same tetrode with PFDs of an angular separation < π/6 radians, and keeping on the highest-rate cell. This selection procedure resulted in a final set of 436 HD cells.

#### Directional and spatial analysis

To determine the firing directions of neurons in global space we projected the 3D nose vector data onto the horizontal plane, binned the firing rate data into 6-deg directional bins and divided the nose vector dwell time in each bin. HD cells were identified based on the model-fitting procedure described below, with a κ (tuning curve sharpness) criterion of 0.5. The resultant cells had tuning curves that looked on visual assessment to be typical HD cells.

#### Model-fitting analysis

Rats moved relatively freely over the hemisphere (see video) but spent less time on the steeper parts. They often tended to circle around the hemisphere, sometimes shuffling around the steeper parts sideways, or else walked down the hemisphere as far as they dared before backing upwards again. This behaviour restricted the full sampling of directions at each location, particularly nearer the equator. This led to our developing an alternative analysis method based on model-fitting to the firing rate profile, as follows.

The FR-direction relationship of a tuning curve is described by a modified von Mises distribution (below).

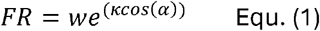

where *FR* is the firing rate, *w* (weight) is a gain factor, κ (kappa) modulates tuning curve sharpness and α (alpha) is the angular distance of the head from the cell’s preferred firing direction, both measured in the plane of interest. We used this relationship to examine the directional firing of neurons over the hemisphere as follows (see Supp. Fig 4c for the pipeline):

1. We used instantaneous firing rates instead of spike counts as these provide a truer picture of the activity of a cell at a given moment. These were derived from individual spike trains at 50 Hz. The elephant electrophysiology toolbox for Python was used ^40^; specifically the function *elephant.statistics.instantaneous_rate*, which implements the Gaussian kernel method described in Shimazaki and Shinomoto ^41^. Optimisation of kernel width, as described in that work, resulted in an extremely large kernel width of 1630ms. Lower kernel widths were judged to be more appropriate, with 200ms decided on as a reasonable compromise between detail and smoothing (Supp. Fig. 2).
2. We next used the recorded data, with directions referenced to the horizontal plane (i.e., the azimuths), to estimate the values of w and k for a given cell, using custom Matlab code (fit_tuning_curves.m). This was done by an iterative model-fitting process, mfit_optimize, ^42^ which searched the parameter space for the best fit. It uses gradient descent to find the best combination of all three parameters, bounding *α* within the range −*π* - *π*, W within the range 0.01 to 100, and κ within the range 0.5-20. We applied this model fit process to the binned tuning curves rather than the raw data because it produced much better fits to the tuning curves and was also very much faster.
3. We based our HD cell selection criterion not on model fit quality, which was often degraded by noise in the data even for obvious HD cells, but rather κ, for which selection above a threshold of 0.5 (the starting value) produced good results according to visual inspection, with evident peaked directional tuning, and Rayleigh vectors almost all > 0.2. This procedure yielded 537 cells. We then identified co-recorded cells that were detected on the same tetrode and had PFDs with < π/6 angular separation, and removed all but the best (highest peak rate) on the basis that they could be the same cell. The final HD cell data set was thus n = 436.
4. We then used the derived parameters to generate a synthetic tuning curve, which was compared with the recorded data using Pearson’s R.
5. The alphas were then transformed according to the four models (details below) using custom-written DAS_transform.m.
6. Using the adjusted alphas, new tuning curve histograms were generated and fitted with von Mises functions using mfit_optimize again using fit_new_all_cells_tuning_curves.m to produce new tuning curve histogram parameters κ, peak rate, Rayleigh vector and model fit (Pearson’s R).
7. Alphas were transformed under GA/DAS/DAH/HA, binned (60 bins, −π to π), and fitted with von Mises functions to obtain κ, w, peak rate, and Rayleigh vector.

### Transformations

The four transformations in step 4 were as follows:

GA rule (gravity axis only): This involved projecting the nose vector onto the horizontal plane and computing azimuth (nose vector in the horizontal plane) in the standard way.

DAS rule (dual axis; surface and gravity): this modulated the alphas according to how much the surface normal axis of the environment local to the rat was rotated around the GA, with the reference (zero) point being in the opposite direction from the cell’s PFD:

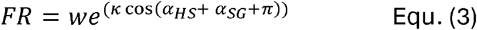

where *α*_*HS*_ is the head rotation around the SA (referenced to the projection of the GA onto the tangent plane) and *α*_*SG*_ is rotation of the SA in the horizontal plane around the GA. The *π* term is added because the zero reference for calculating head α is taken to be when maximal firing (local PFD) coincides with the rat facing upwards, which occurs at the opposite side of the sphere from the cell’s global PFD.

DAH rule (dual axis; head and gravity): this modulated the alphas according to how much the DV axis of the head was rotated around the GA, with, again, the reference (zero) point being in the opposite direction from the cell’s PFD. The modified von Mises equation describing the head-referenced dual-axis rule (DAH) is:

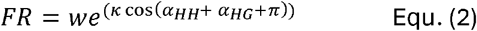

Where *α*_*HH*_ is the alpha of the head rotation in the head plane around the HA (referenced to the projection of the GA onto the head plane) and *α*_*HG*_ is rotation of the HA in the horizontal plane around the GA.

HA rule (head axis only): This ignores the tilt of the environment surface or the head and simply updates according to left-right turns around the head axis, ignoring the head axis azimuth. We expected this to produce poor model fits and this manipulation was included as a null control.

#### Elevation analysis

We defined elevation as the angle of the nose vector relative to the horizontal plane. Since HD cells sensitive to this parameter have been reported previously ^10^, we investigated the distribution of elevation angles, from -π/2 (nose pointing down) to +π/2 (nose pointing up).

### Statistics and reproducibility

Sample size was determined based on prior similar experiments on HD cells over a 30-year period. A significance level of p < 0.05 was applied.

Given that cluster-cutting is necessarily imperfect, and to ensure we did not include a given cell more than once in the data set, we applied a conservative acceptance criterion such that two simultaneously recorded “cells” needed to have tuning curves with PFDs > π/6 radians apart to be both included (otherwise we included only the highest-rate one).

Because analysis was at the cell level but the cells were nested within animals, we used a linear mixed-effects model to analyze the tuning curve parameters. To compare the four models (GA, DAS, DAH and HA) for each dependent measure (peak rate, Rayleigh vector length, κ, and model-fit metric), we fit a model of the form:

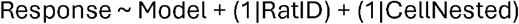

where Model had four levels (GA, DAS, DAH, HA), and CellNested indexed cell identity nested within rat. Models were fit by maximum likelihood. We report the omnibus fixed effect of Model and planned contrasts testing the a priori hypotheses that DAH > DAS, GA, HA (Family A, Holm-adjusted) and that DAS > GA (Family B).

## Data availability

The datasets analyzed during the current study, including those subsets used to generate the figures, are available from https://doi.org/10.6084/m9.figshare.31222660

## Code availability

Code used for this article is available from

https://doi.org/10.6084/m9.figshare.31222660

## Acknowledgements

The work was supported by a BBSRC training grant to MW (BB/M009513/1), a Wellcome Principal Research Fellowship to NB (202805/Z/16/Z) and a Wellcome Investigator Award to KJ (103896/Z/14/Z).

For the purpose of Open Access, the author has applied a Creative Commons Attribution (CC BY) license to any Author Accepted Manuscript version arising from this submission.

## Author contributions

MW gained funding, designed the experiment, designed and built the apparatus, collected the data, conceptualized and performed analyses, edited the manuscript; JS provided advice and support, and edited the manuscript; NB provided advice and support, and edited the manuscript; KJ conceptualized the experiment, gained funding, conceptualized and performed analyses, and edited the manuscript.

