## Supplementary tables and figures for "Head direction cells use a head-referenced dual-axis updating rule in 3D space"

### Supplementary information

#### Supplementary Tables

###### Supplementary Table 1

|  |  | **ADN** | | **POS** | | **Total** | |
| --- | --- | --- | --- | --- | --- | --- | --- |
|  |  | *Mean* | *s.e.m* | *Mean* | *s.e.m* | *Mean* | *s.e.m* |
| **GA** | Pk rate (Hz) | 7.89 | 0.41 | 3.62 | 0.36 | 6.51 | 0.32 |
|  | R vector | 0.45 | 0.01 | 0.50 | 0.02 | 0.47 | 0.01 |
|  | kappa | 1.73 | 0.10 | 3.04 | 0.27 | 2.15 | 0.12 |
|  | Model fit | 0.37 | 0.01 | 0.33 | 0.01 | 0.36 | 0.01 |
| **DAS** | Pk rate (Hz) | 8.68 | 0.47 | 3.74 | 0.38 | 7.08 | 0.36 |
|  | R vector | 0.48 | 0.01 | 0.52 | 0.02 | 0.49 | 0.01 |
|  | kappa | 1.88 | 0.10 | 3.16 | 0.28 | 2.30 | 0.12 |
|  | Model fit | 0.38 | 0.01 | 0.34 | 0.01 | 0.37 | 0.01 |
| **DAH** | Pk rate (Hz) | 9.94 | 0.53 | 4.24 | 0.40 | 8.10 | 0.40 |
|  | R vector | 0.53 | 0.01 | 0.56 | 0.02 | 0.54 | 0.01 |
|  | kappa | 2.94 | 0.15 | 4.88 | 0.37 | 3.57 | 0.16 |
|  | Model fit | 0.47 | 0.01 | 0.39 | 0.01 | 0.44 | 0.01 |
| **HA** | Pk rate (Hz) | 5.79 | 0.30 | 2.14 | 0.20 | 4.61 | 0.23 |
|  | R vector | 0.18 | 0.01 | 0.19 | 0.01 | 0.18 | 0.01 |
|  | kappa | 0.73 | 0.10 | 0.58 | 0.02 | 0.68 | 0.07 |
|  | Model fit | 0.00 | 0.00 | 0.01 | 0.00 | 0.00 | 0.00 |

**Supplementary Table 1:** Tuning curve parameters in standard horizontal coordinates (GA) and under the three transformations (DAS, DAH and HA), compared between brain regions

###### Supplementary Table 2

|  | **GA** | **DAS** | **DAH** | **HA** |
| --- | --- | --- | --- | --- |
| **Pk rate** | 26 | 78 | 280 | 52 |
| **Best R vector** | 44 | 42 | 343 | 7 |
| **Best kappa** | 17 | 41 | 367 | 11 |
| **Best model fit** | 22 | 32 | 382 | 0 |

**Supplementary Table 2:** No. cells for which a given model yielded the best value. The DAH model yielded the best value for many more cells than the other models. See text for statistical comparisons.

###### Supplementary Table 3

| **Variable** | **Comparison** | **Contrast** | ***T***  **(one-tailed)** | ***p*** | **Effect size *(d)*** |
| --- | --- | --- | --- | --- | --- |
| Peak rate | DAH vs DAS | 1.01 | 6.04 | <0.0000 | 0.41 |
|  | DAH vs GA | 1.59 | 9.46 | <0.0000 | 0.64 |
|  | DAH vs HA | 3.49 | 20.79 | <0.0000 | 1.41 |
|  | DAS vs GA | 0.57 | 3.42 | <0.0000 | 0.23 |
| R. vector | DAH vs DAS | 0.05 | 7.52 | <0.0000 | 0.51 |
|  | DAH vs GA | 0.08 | 11.23 | <0.0000 | 0.76 |
|  | DAH vs HA | 0.36 | 53.34 | <0.0000 | 3.63 |
|  | DAS vs GA | 0.02 | 3.72 | 0.0001 | 0.25 |
| kappa | DAH vs DAS | 1.27 | 10.76 | <0.0000 | 0.73 |
|  | DAH vs GA | 1.42 | 11.97 | <0.0000 | 0.81 |
|  | DAH vs HA | 2.89 | 24.37 | <0.0000 | 1.65 |
|  | DAS vs GA | 0.14 | 1.20 | 0.12 NS | 0.08 |
| Model fit (Pearson’s R) | DAH vs DAS | 0.08 | 10.96 | <0.0000 | 0.74 |
|  | DAH vs GA | 0.08 | 12.26 | <0.0000 | 0.83 |
|  | DAH vs HA | 0.44 | 64.34 | <0.0000 | 4.36 |
|  | DAS vs GA | 0.01 | 1.32 | 0.09 NS | 0.09 |

**Supplementary Table 3:** Linear mixed model analyses

#### Supplementary Figures

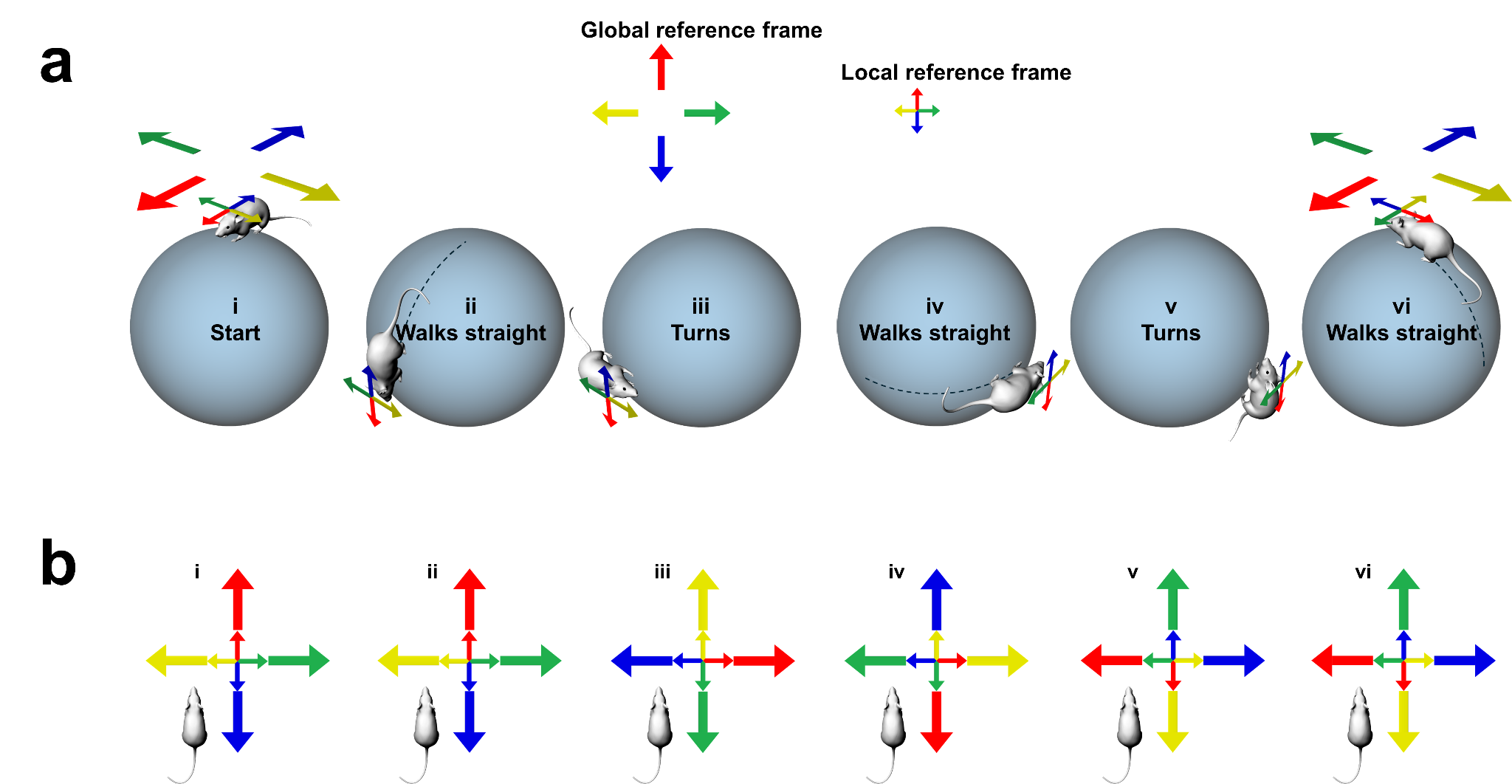

**Supplementary Figure 1: Berry-Hannay errors following exploration in 3D**. (a) A rat travels over a sphere, carrying its internal, local reference frame (small colored arrows) with it. By the end of the journey, having returned back to a horizontal alignment, its local reference frame has dissociated from the global one (large colored arrows). (b) The coordinate reference frames from (a) expressed relative to the rat (egocentrically). The first turn (iii) rotates the rat in its internal coordinate frame (it is now facing the “yellow” direction; inner arrows) but the local frame remains aligned to the global frame (outer arrows). The second turn (v) both rotates the rat within its own frame *and* the local frame in the global frame, the result being a dissociation. The dissociation occurs because in 3D, rotation can indirectly occur in a given plane by rotations occurring in the orthogonal planes, which a simple planar ring attractor would not detect.

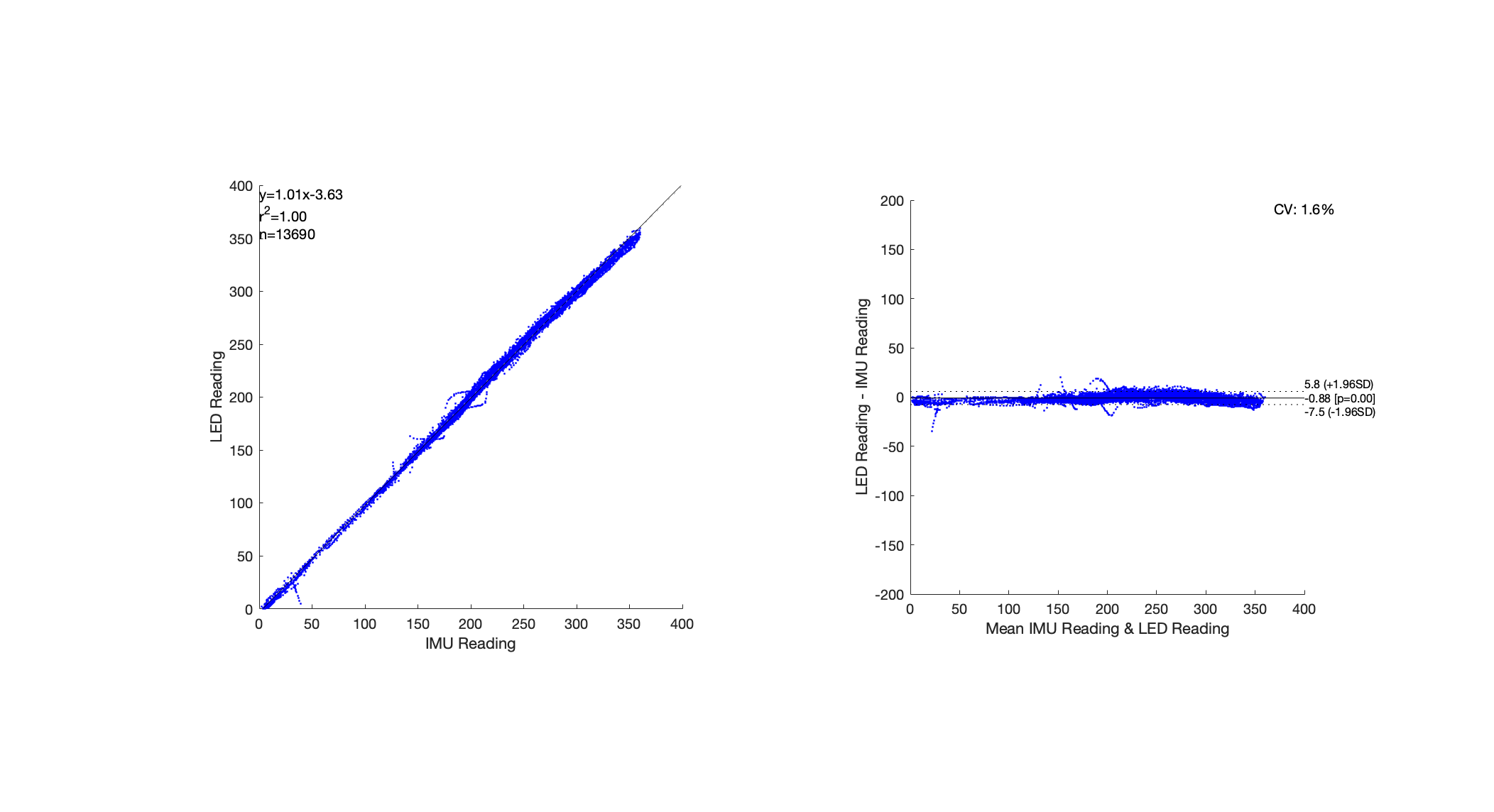

**Supplementary Figure 2: IMU and classic LED-based head direction recording methods show a high degree of agreement**. Bland-Altman analysis of paired recording with LEDs and IMU: blue points are individual readings at 50Hz. Left shows LED and IMU readings plotted against one another, with line of best fit, R^2^ value, and number of samples. Right shows mean difference between the two sets of readings.

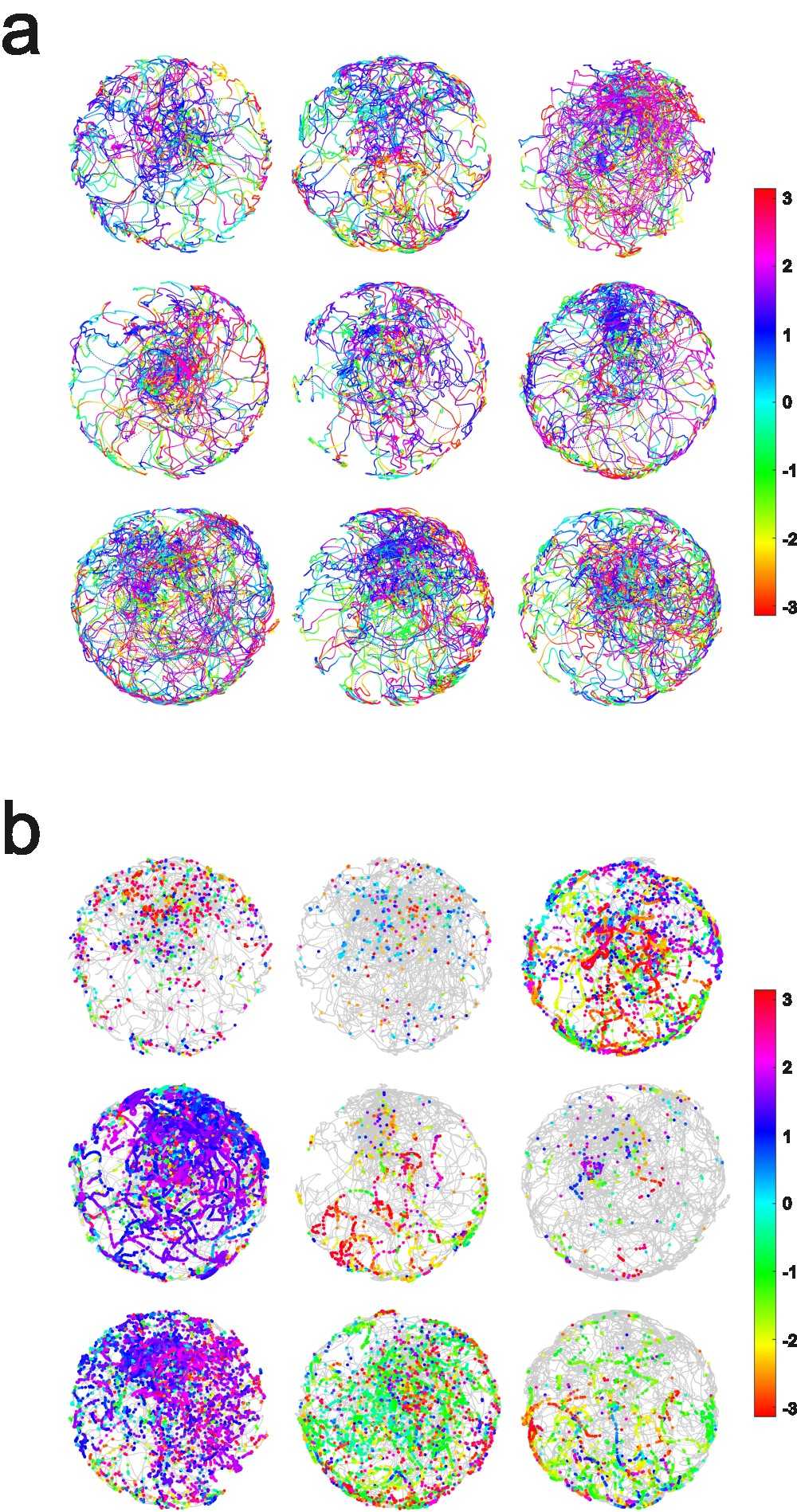

**Supplementary Figure 3: Distribution of position and facing direction**. (A) Example sessions (every 12^th^ session) to show typical distributions of location and facing direction as animals traversed the sphere. Although they spent more time at the top, they did frequently walk down the side to the equator. Note the inhomogeneous distribution of facing directions (as indicated by the key, showing direction in radians). (B) Nine example cell recording trials (every 48^th^ cell) showing the location and azimuth of each spike, color-coded as in A.

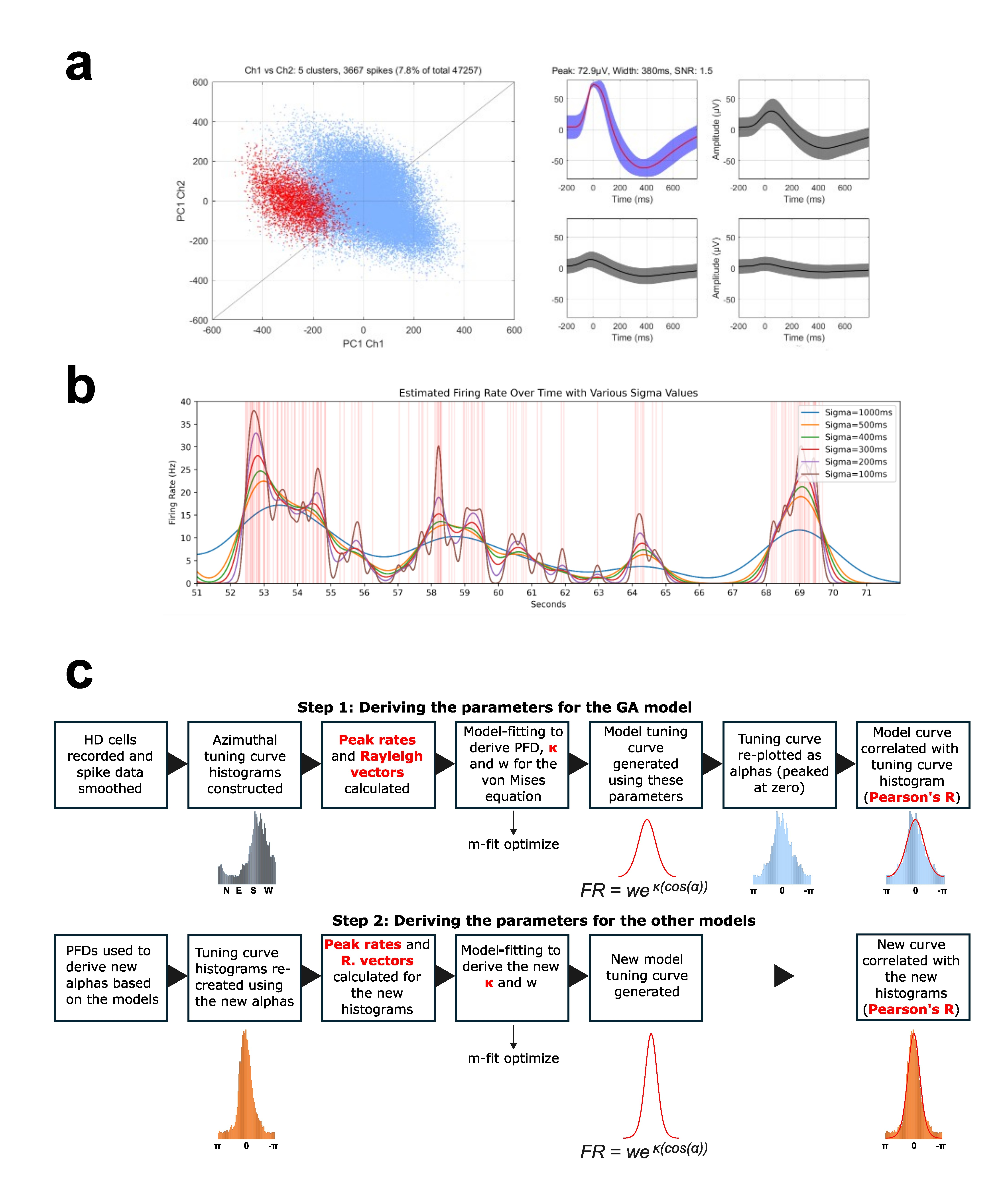

**Supplementary Figure 4:** **Neuronal analyses** (a) Left: cluster space for a hemisphere session, showing a HD cell in red, other units in blue. Right, waveforms for the same cell, highest amplitude waveform in red. Plots show waveform means and standard deviations. Peak, width, and signal to noise ratio (SNR) also shown. (b) Gaussian smoothing to generate the FR profile. Example spiketrain (red vertical lines) and estimated instantaneous firing rates calculated using gaussian kernels (Shimazaki and Shinomoto, 2010). Various sigma values (kernel widths) are shown in different colors. Sigma value used in final analysis was 200ms (purple). (c) Analysis pipeline for the tuning curves. The original tuning curve histograms were converted from global coordinates (N = North, E = East etc) to PFD-centered coordinates (alphas). The histograms were then modeled by a modified von Mises function, and correlated with the tuning curve histograms using Pearson’s R. The procedure was then re-done using the re-valued alphas, based on the algorithms for the three different models (DAS, DAH and HA). The four parameters for comparing model quality are shown in bold red text. The Pearson’s R model fit was taken as the measure of the best model.

**Supplementary Figure 5: Example tuning curves plotted according to the four models.**  Forty cells are shown (approx. every 10^th^ cell in the data set; x axis is nose vector in the relevant reference frame; y axis shows firing rate in Hz) . The best model for each cell (defined by the best Pearson’s R model fit) is indicated in the header and color code for each cell. The great majority of tuning curves were sharpest and tallest under the DAH model.

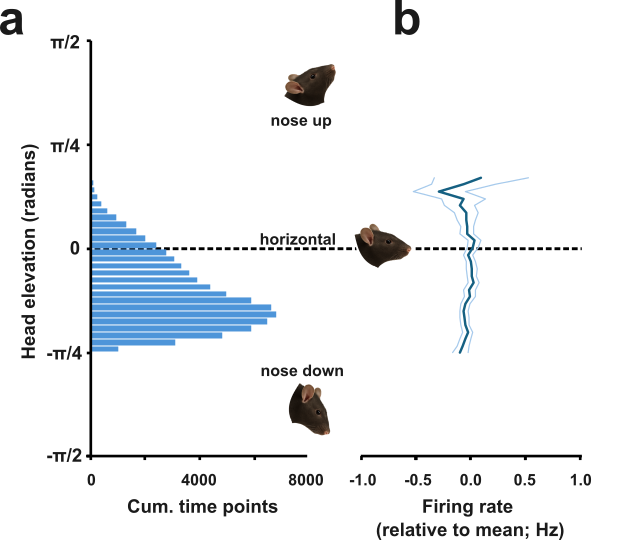

**Supplementary Figure 6: Calculation of elevation tuning.** (a) Distribution of nose elevation angles. (b) Firing rates relative to the cell’s mean, for all 2270 cells as a function of nose elevation (mean +/- s.e.m)

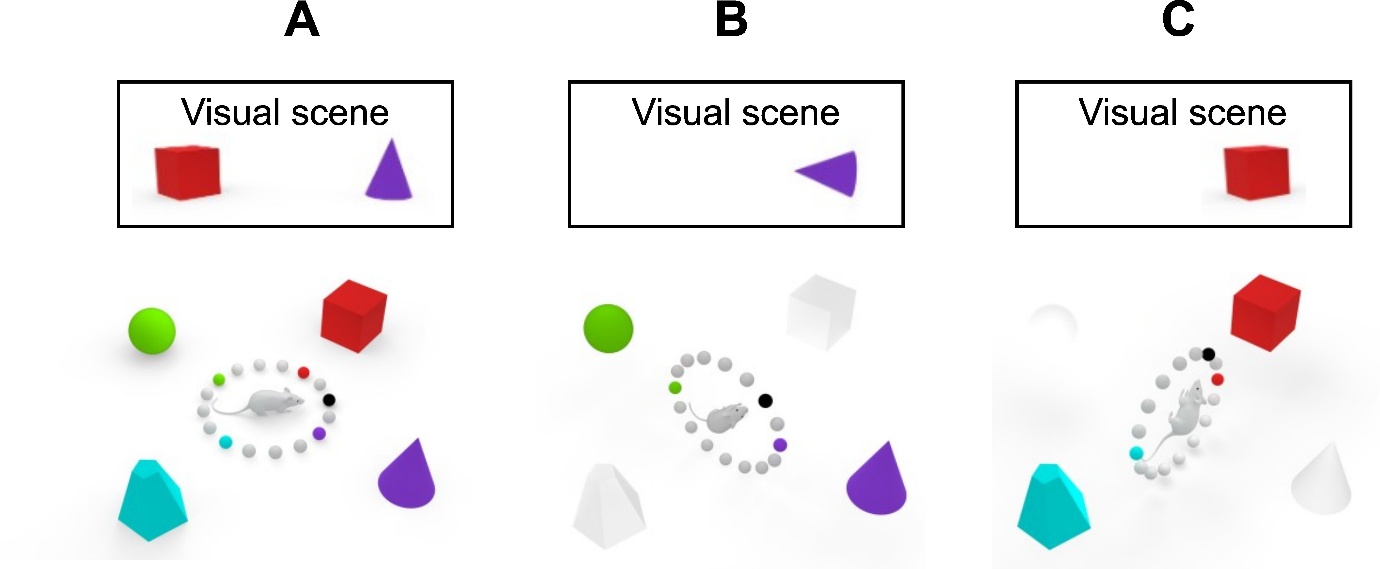

**Supplementary Figure 7:** Landmark-setting of the ring attractor in 3D. (A) On the horizontal plane, landmarks drive different parts of the ring attractor so as to anchor network activity to the environment. The black sphere represents the currently active neuron in this schematic; the colored spheres represent the neurons that would be active if the animal faced towards the correspondingly color-coded landmark. The inset shows the visual scene from the animal’s perspective: this scene has been linked via learning to a particular part of the ring attractor. (B) In a vertical pose the same landmarks could be used to drive the same parts of the ring attractor, although some will be obscured (greyed-out landmarks) and the orientation of the landmark is altered. (C) In the orthogonal vertical plane, the same logic holds: the visual scene, with obscured or rotated landmarks, can still drive the same part of the ring attractor and help to set the network activity appropriately. Note that static vestibular cues to pose, from the otoliths, may also in principle contribute a drive to the attractor.

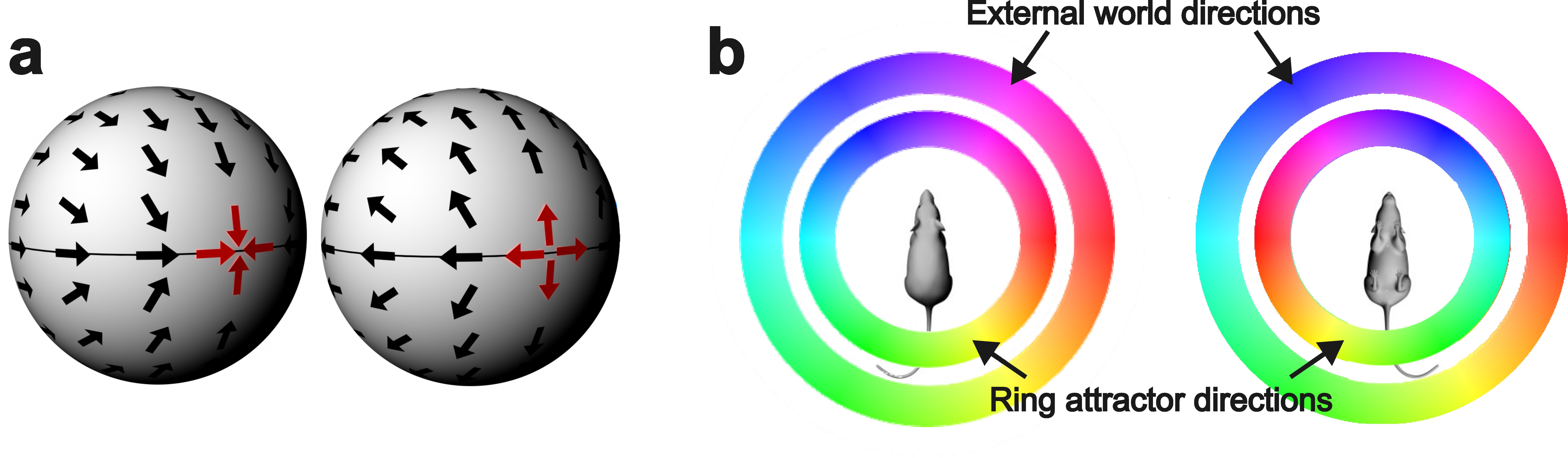

**Supplementary Figure 8. Limitations of planar encoding in 3D space.** (a) The problem with the DA rule, which yields a planar signal, is the inevitability in the HD cell vector field of singularities (at least two, at the “front” and “back” of the sphere), where the vector abruptly needs to change direction. This is described mathematically by the Hairy Ball theorem. (b) Also, when the animal inverts, the relationship between directions in the local ring attractor and directions in the global external environment reverses.

#### Supplementary movie

See <https://doi.org/10.6084/m9.figshare.29649560>
